# Physiological characterization of single gene lysis proteins

**DOI:** 10.1101/2023.10.16.562596

**Authors:** S. Francesca Antillon, Thomas G. Bernhardt, Karthik Chamakura, Ry Young

**Affiliations:** Department of Biochemistry and Biophysics, Texas A&M University, College Station, TX 77843-2128, United States; Center for Phage Technology, Texas A&M AgriLife Research, College Station, TX, 77843-2128, United States; Department of Microbiology, Blavatnik Institute, Harvard Medical School, Boston, MA 02115; HHMI, Chevy Chase, MD 20815

## Abstract

Until recently only 11 distinct Sgls (<u>si</u>ngle <u>g</u>ene lysis proteins) have been experimentally identified. Of these, three have been shown to be specific inhibitors of different steps in the pathway that supplies Lipid II to the peptidoglycan (PG) biosynthesis machinery: Qβ A_2_ inhibits MurA, ϕX174 E inhibits MraY, and Lys from coliphage M inhibits MurJ. These Sgls have been called “protein antibiotics” because the lytic event is a septal catastrophe indistinguishable from that caused by cell wall antibiotics. Here we propose to designate these as members of type I Sgls, to distinguish them from another Sgl, the L protein of the paradigm ssRNA phage MS2. Although none of the other distinct Sgls have significant sequence similarity to L, alignments suggested the presence of four domains distinguished by hydrophobic and polar character. The simplest notion is that these other Sgls have the same autolytic mechanism and, based on this, constitute type II.

Although the number of experimentally confirmed Sgls has not changed, recent environmental metagenomes and metatranscriptomes have revealed thousands of new ssRNA phage genomes, each of which presumably has at least one Sgl gene. Here we report on methods to distinguish type I and type II Sgls. Using phase-contrast microscopy, we show that both classes of Sgls cause the formation of blebs prior to lysis, but the location of the blebs differs significantly. In addition, we show that L and other type II Sgls do not inhibit net synthesis of PG, as measured by incorporation of ^3^[H]-diaminopimelic acid. Finally, we provide support for the unexpected finding by Adler and colleagues that the Sgl from Pseudomonas phage PP7 is a type I Sgl, as determined by the two methods. This shows that the sharing the putative 4-domain structure suggested for L is not a reliable discriminator for operational characterization of Sgls. Overall, this study establishes new ways to rapidly classify novel Sgls and thus may facilitate the identification of new cell envelope targets that will help generate new antibiotics.

## Introduction

For host lysis, double-strand DNA phages encode multiple lysis proteins that actively attack each layer of the envelope according to a tightly-regulated temporal schedule (1, 2). More than a dozen different functional types of proteins have been linked to this “multi-gene lysis” (MGL) process, with some phages encoding as many as six. By contrast, small lytic phages with single-strand RNA and DNA chromosomes lack the genomic space for an MGL system. Instead, these phages, the *Microviridae* and the ssRNA *Leviviricetes*, liberate their progeny by inducing lysis through expression of a single gene (3, 4). These genes have been collectively named *sgl* (single gene lysis; product Sgl). Importantly, Sgls do not have bacteriolytic enzyme activities and thus each is, formally, an inducer of host autolysis. As such, Sgls are of interest in terms of development of new antibiotic strategies, each representing a potential modality for instigating autolysis of a bacterial cell (5). The position and character of the *sgl*s are particularly striking in the ssRNA phages, which have three core genes that encode virion and replication proteins (Fig. 1A). In contrast to the core genes, which are always in the same order and approximately the same size, the location and size of the *sgl* is highly variable, embedded out of frame in one of the three core genes or positioned in an intergenic space (3). This strongly indicates that there is an independent evolutionary path for each *sgl*. The classic case is the first Sgl, identified by genetic analysis in the canonical F-specific coliphage MS2 (6). In this case, Sgl^MS2^, given the gene name *L* for lysis, is a 75-codon reading frame overlapping the *coat*-*rep* junction (Fig. 1). Its Sgl character was unambiguously demonstrated by its ability to cause autolysis by induction from plasmid expression vectors (7-9). Until recently the Sgls of only 9 other plaque-forming phages have been identified by the same functional criterion (3). This includes two Sgls that have been extensively studied: E, the Sgl of the canonical ssDNA Microvirus ϕX174, and A_2_, the dual function maturation protein/Sgl of Qβ, another well-studied F-specific ssRNA phage of *Escherichia coli*. In addition, Sgls have been identified for ssRNA phages specific for the pili of conjugative plasmids of *E. coli* (HgaI, C-1, and M) and *Pseudomonas* (PRR1), as well for the polar retractable pili of *Pseudomonas* (PP7), *Acinetobacter* (AP205), and *Caulobacter crescentus* (phiCB5). Definitive mechanistic information is available for only E and A_2_. Biochemical and structural evidence indicates that each acts by binding and inhibiting an essential enzyme in the pathway for biosynthesis of Lipid II, the precursor of peptidoglycan biosynthesis: MraY and MurA, respectively (10-13). In addition, genetic and molecular evidence has indicated that the Sgls of coliphage M and the *Pseudomonas* phage PP7 inhibit MurJ, which flips Lipid II to the periplasm as the last step of the pathway. The sobriquet “protein antibiotic”, originally coined for A_2_, alluding to the functional similarity to beta-lactams and other cell wall antibiotics (14), could reasonably be extended to all four of these Sgls.

**Figure 1.**
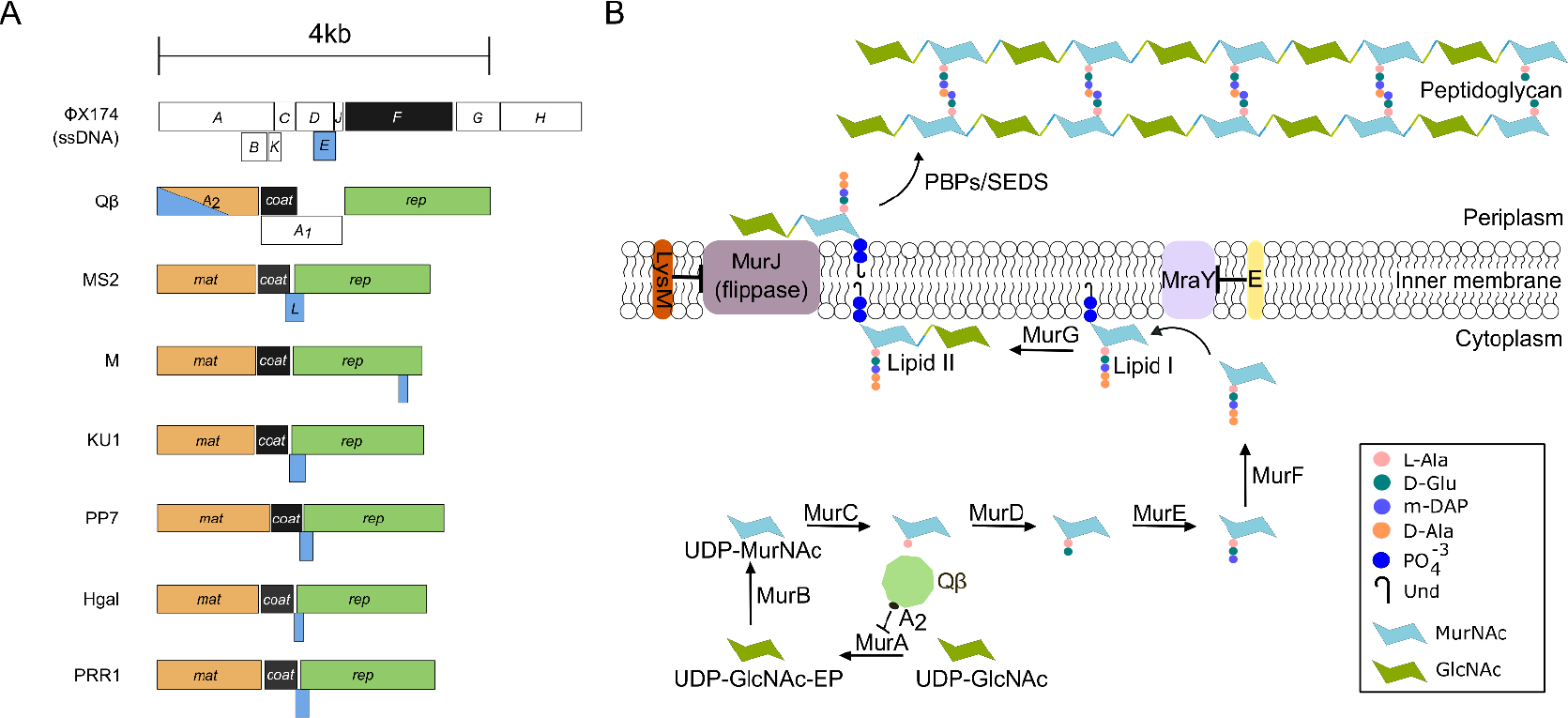
Peptidoglycan biosynthesis pathway and Sgl inhibition. **A**. Genomes of ssDNA (ϕX174) and ssRNA (Qβ, MS2, M, KU1, PP7, Hgal, and PRR1) phages. Adopted from (9). SsRNA phages have three core conserved genes in the same order: *mat* (orange), *coat* (black), and *rep* (green). These phages also encode a lysis gene (blue) that is embedded in an alternative reading frame in one of the three core genes or in intergenic spaces. In Qβ the maturation gene has a dual function, serving as both the maturation protein and the Sgl. ***B***. Sgls of ssRNA and ssDNA phages such as ϕx174,Qβ, and M inhibit different enzymatic steps in the PG biosynthesis pathway.

In contrast, mechanistic information for the other six Sgls is sparse. MS2 L has been the subject of multiple studies, beginning in the mid-1980s with a series of collaborative works from the groups of J. van Duin and J. Höltje. One monograph reported pulse-labelling studies showing that L did not affect the rate of PG biosynthesis (15). In addition, immuno-electron microscopy data were cited indicating that L was enriched in zones of adhesion between the IM and OM (16). Finally, a synthetic peptide corresponding to the C-terminal 25 AA of L was reported to have pore-forming character for liposomes, leading to the model that L-mediated autolysis involved membrane pore formation (17). More recently, Mezhyrova et al. (18) used in vitro liposome permeabilization experiments and in vivo cryo-electron microscopy to support this model. In addition, our group reported a genetic analysis of L (8). The results and comparison with L sequences from closely related F-specific MS2-like phages suggest that L has four domains, including an essential hydrophobic domain preceding a conserved Leu-Ser dipeptide sequence (Supp. Fig. 1). Moreover, the highly cationic N-terminal domain, although previously shown to be non-essential for lytic function, was found to confer dependence of the host chaperone DnaJ (7, 9). Other than the three Sgls shown to be inhibitors of the lipid II biosynthesis pathway, all the other Sgls have similar domains, including the LS dipeptide, although none have significant sequence similarity (Supp. Fig 1) (8). Taken together with the studies on L, these findings suggested that L and the other 5 Sgls might be functionally related (“L-like”). Recently the significance of the LS dipeptide was brought into question when Sgl^PP7^, which has a similar multi-domain structure, was implicated as being an inhibitor of MurJ by multi-copy suppression and genetic studies (19).

Recently thousands of new ssRNA phage genomes have been discovered in environmental transcriptomes and viromes (20-23). The diversity of this new ssRNA genomic space is staggering but not surprising, considering the high mutagenesis rate that characterizes RNA-dependent RNA polymerases (24). Importantly, homologs of the known Sgls were very rare in these new genomes. Moreover, 33 new sgl genes were found by bioinformatic and functional analysis, mapping to at least 18 different positions in the genome (19). This unparalleled diversity raises the alluring possibility that Sgl lysis targets might titrate many of the known steps of PG biosynthesis and homeostasis, plus new ones, leading to multiple new antibiotic targets. Some success at identifying and characterizing the *sgl* genes from these collections was obtained by synthesizing more than 300 potential *sgl* genes, selected for having a potential transmembrane domain (TMD) as well as plausible reading frames and translational start (20). When these *sgl* candidates were synthesized and tested for inducible lysis in *E. coli*, about 10% were shown to have lytic function, a reasonable success rate considering that the bacterial hosts for these hypothetical phages are not known. Given the number of Sgls to be tested, it would be very useful to be able to segregate the “protein antibiotic” Sgls from the L-like Sgls, based on the notion that each identifies a sensitive molecular interaction surface in a potential antibiotic target.

Given this new wide perspective and the likelihood that there will be many fundamentally different mechanistic systems, we propose to organize the Sgl database into a series of types, as has been done profitably with the CRISPR system. We propose to reclassify the Sgls that inhibit PG biosynthesis as type I Sgls and the L-like Sgls as type II. For the purposes of the rest of this report, we will use this terminology. In addition, we will follow historical precedent in referring to the well-studied Sgls from MS2, Qβ, and ϕ174 as L, A_2_ and E, respectively but all others as Sgl^X^, where X is the name of the phage.

## Results

### Type I Sgls but not Type II Sgls inhibit net PG synthesis

As noted above, it has been reported that, for lytic inductions of *L*, the rate of PG synthesis, as measured by pulse-labeling with ^3^[H]mDAP, is unaffected up to the time of lysis (15). In contrast, for both E and A_2_, net PG synthesis, as measured by continuous incorporation of the same label, is halted at least 20 min before the lytic event (14, 25). These results raised the possibility that type I Sgls can be distinguished from type II using PG labeling. The simplest method is to follow the incorporation of ^3^[H]mDAP into material that is insoluble in boiling SDS; i.e., the sacculi. However, ^3^[H]mDAP labeling requires minimal medium, which is done with glucose as the carbon source. Moreover, in our hands, reproducible results with cloned *sgl* genes require the use of tightly regulated arabinose (Ara)-inducible vectors (see Supplementary Fig. 2), because appreciable basal level expression of these highly toxic genes leads to growth anomalies prior to induction. However, since Ara inductions are subject to catabolite repression, a different carbon source was required. Among the available carbon sources that do not cause catabolite repression, studies with *E. coli* B/r have shown that glycerol supports reasonable growth rates if the minimal medium is supplemented with all the amino acids (26-28). We tested glycerol as a carbon source for our standard induction system, using a *lysA* derivative of the *E. coli* K-12 strain MG1655 (Fig. 2a,b). Cultures exhibited reproducible growth rates of ~1 doubling/hr and, depending on the cloned *sgl* gene, showed robust, distinct lysis profiles at 10 to 30 min after addition of arabinose. In the case of two well-established type I Sgls, E and Sgl^M^, net incorporation of ^3^[H]mDAP stopped abruptly long before lysis was apparent (Fig. 3). In contrast, incorporation continued unabated until lysis in the case of the type II paradigm, L, consistent with the previous pulse-labelling results (15). Similar results were obtained with the Sgls from phages KU1 and PRR1, indicating that both are Type II Sgls (Fig. 3). A modified labeling system was required for the *sgl* of phage Hgal, which for unknown reasons failed to support lysis when induced from the Ara-inducible platform (Supp. Fig. 2). In this case, the crystal violet (CV)-inducible vector pJexD was used (Supp. Fig 3) and again, labeling continued until lysis (Fig. 3). Taken together these data indicate all three are type II Sgls. Although none of these genes have been experimentally characterized, they all have marginal sequence similarity to MS2 L, have a four-domain primary structure, and are encoded in a reading frame similarly positioned to *L* overlapping the *coat* – *rep* boundary (8).

**Figure 2.**
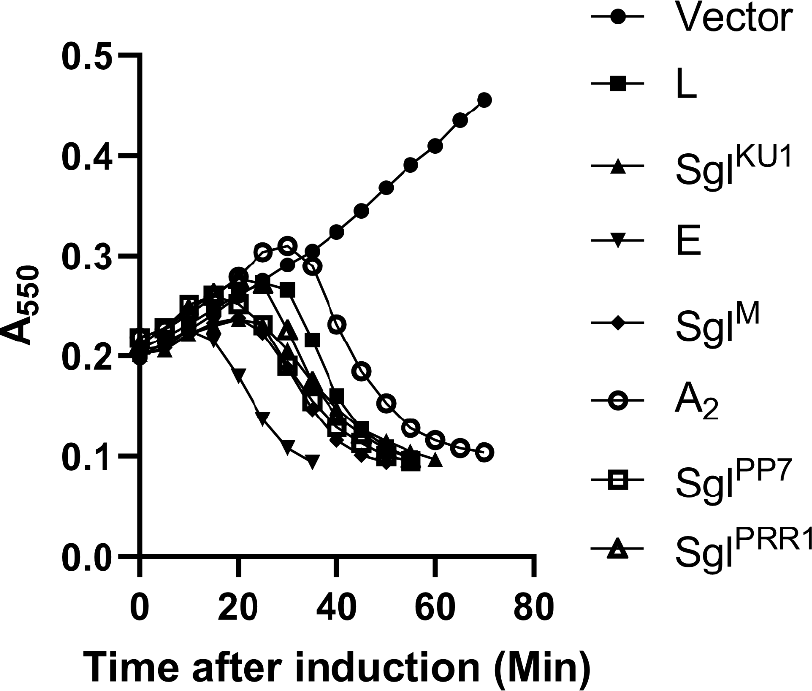
Sgl lysis profiles in defined minimal media. Lysis profiles of Sgls in minimal media supplemented with 16 amino acids, 0.01% thiamine, and 0.2% glycerol as the carbon source. When the cultures reached an OD of ~0.2 they were induced with 0.4% L-arabinose. From top to bottom: pBAD24 Vector with no Sgl (closed circles), L (closed squares), Sgl^KU1^ (closed upward triangles), E (closed downward triangle), Sgl^M^ (closed diamonds), A_2_ (open circles), Sgl^PP7^ (open squares), and Sgl^PRR1^ (open triangles).

**Figure 3.**
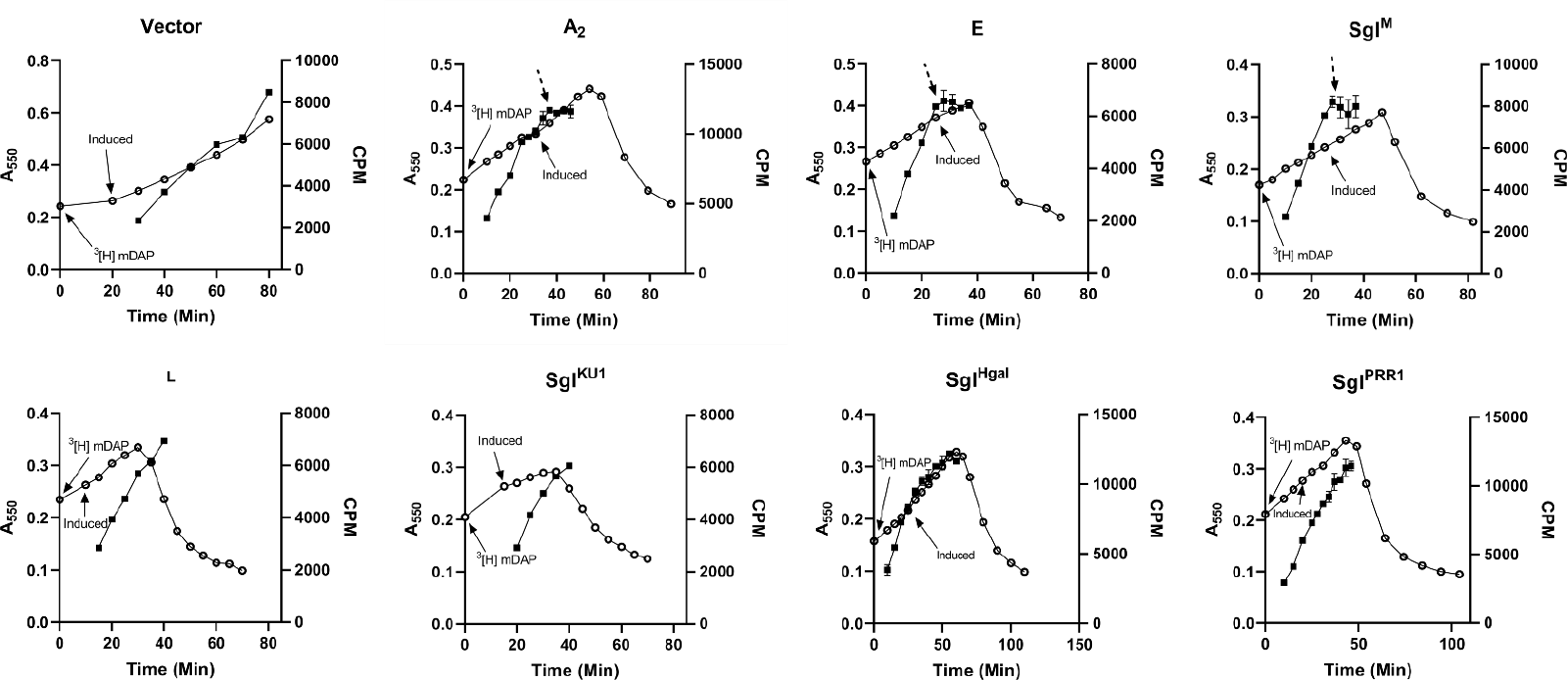
^3^[H] mDAP labeling of peptidoglycan. Growth curve of *E. coli* is shown as open circle (⭘) and the amount of ^3^[H] mDAP counted at different time points is shown as closed squares (■). The first arrow indicates when ^3^[H] mDAP was added to the bacterial culture and the second arrow indicates the induction of the plasmid. The dashed arrows represent the time at which incorporation of ^3^[H] mDAP CPM ceased. Each construct has a minimum of one repeat and the representative one is shown in each case. For type I Sgls, after induction, each time point was filtered twice (see methods). The average is plotted, and the error bars represent the variation.

### Type I and Type II Sgls cause different lytic morphology

Exposure to cell wall antibiotics causes a distinct morphological change in growing bacteria (29, 30). Most cells undergo catastrophic septal failures, with a large bleb emanating from the mid-cell. This presumably reflects the requirement for massive de novo PG biosynthesis localized to the developing septum, compared to the highly distributed nature of PG biosynthesis in the elongating cell walls (31). In principle, all type I Sgls should cause the identical lytic morphology, given the common end-point (i.e., starvation for periplasmic Lipid II. Indeed, *E. coli* cells infected with the ssDNA phage ϕX174, which uses the canonical type I Sgl E, exhibits mid-cell blebbing (32). This lysis morphology has been recapitulated with the cloned *E* gene (33-35), demonstrating that expression of E is necessary and sufficient not only for lysis but also for the particular lysis morphology characterized by a septal catastrophe. We tested if this morphological phenotype could be used as a criterion for distinguishing type I and type II Sgls. We reproduced the E finding and found that the septal blebbing was also characteristic of other type I Sgls, A_2_ and Sgl^M^ (Fig. 4,5,6). In contrast, L and the putative type II Sgls of KU1, PPR1, and Hgal gave rise to more distributed, numerous and smaller blebs (Fig. 4,5). Moreover, the blebs exhibited by the type II Sgls were significantly less stable, as judged by the frequency of lysed cells in microscopy fields (Sup. Fig. 4). Indeed, being able to visualize any significant number of intact blebs, as judged by phase contrast, required the addition of Mg^++^ ions and sucrose to stabilize the envelope. Stabilizing conditions were used to image both type I and type II Sgls for a direct comparison.

**Figure 4.**
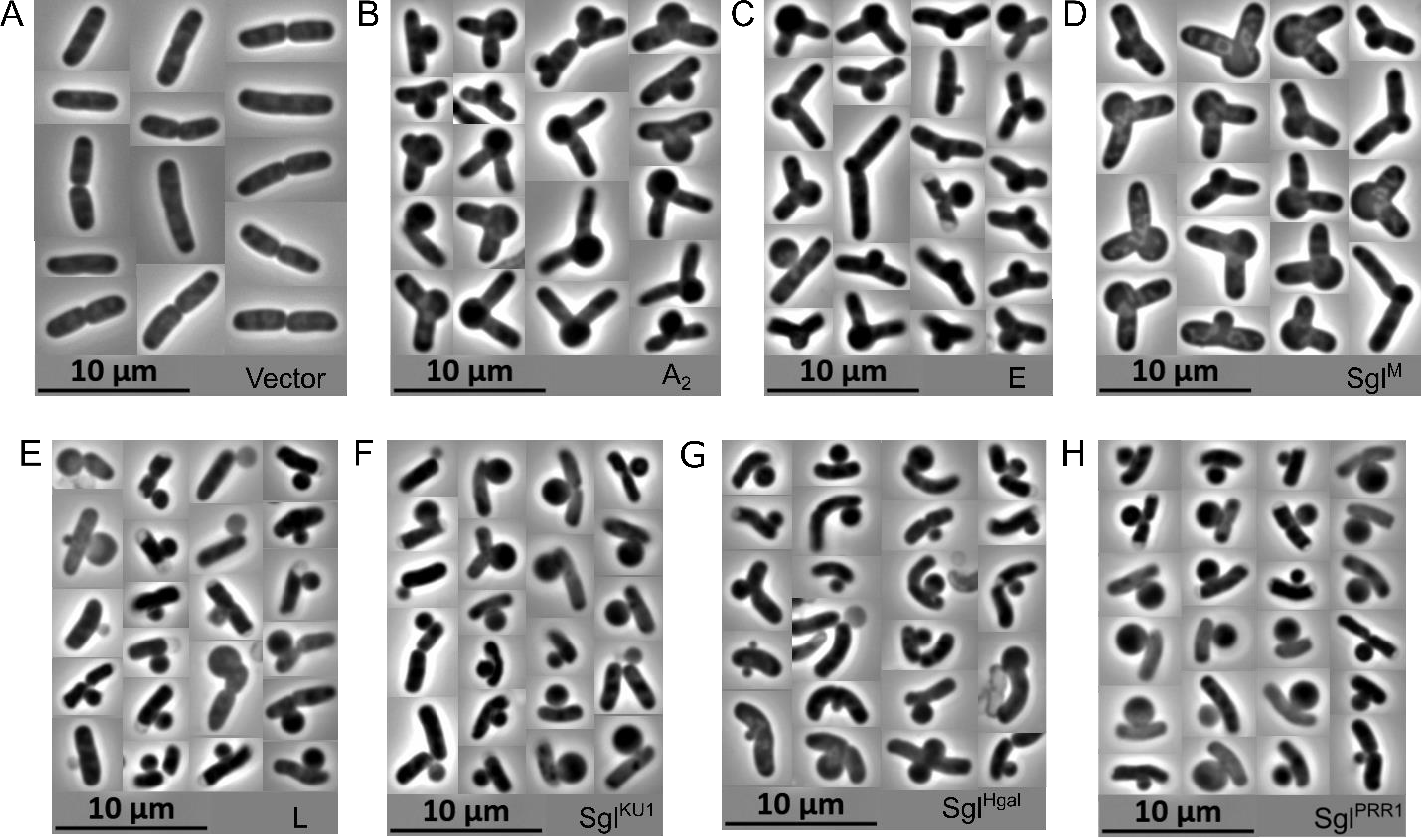
Type I and type II lysis morphology. Shown are phase-contrast images of *E. coli* after induction of different lysis genes. **A**. Vector only **B**. A_2_ **C**. E **D**. Sgl^M^ **E**. L **F**. Sgl^KU1^ **G**. Sgl^Hgal^ **H**. Sgl^PRR1^. For panels A-G, cultures were grown at 37°C to an optical density of ~0.2 and induced with 0.4% L-arabinose. For panel H, cultures were induced with 0.075µM crystal violet. Ten minutes prior to lysis, 1.5μL samples were taken from culture flasks, put onto glass slides, covered with a cover slip, and imaged using the 100X objective lens. Collages were made using Inkscape and each represents the population of cells seen in triplicate imaging experiments. Furthermore, to minimize bias in the selection of cells with blebs for the collages, all cells showing blebs on an image were incorporated into collages.

**Figure 5.**
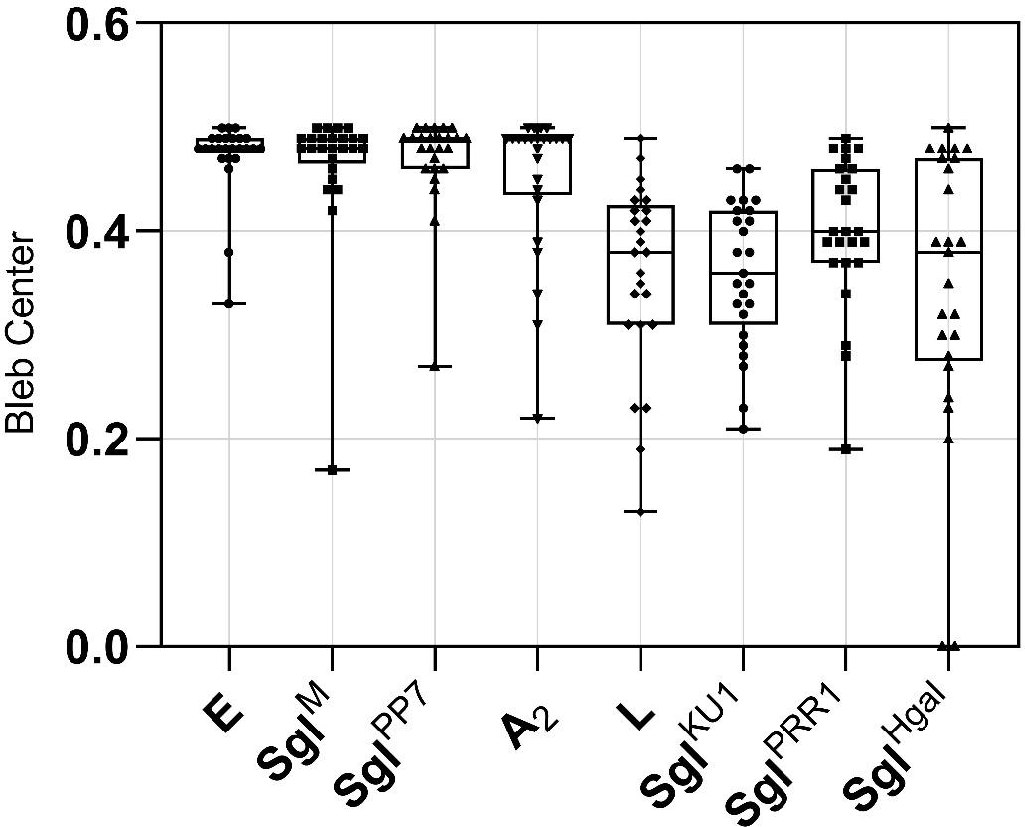
Analysis of lesions caused by Sgls. The y-axis represents the location of the bleb along the cell length with 0.0 representing the poles of the cell and 0.5 representing middle of the cell. X-axis indicates the type of Sgl each set of measurements, including type I (E, Sgl^M^, Sg^PP7^, and A_2_) and type II (L, Sgl^KU1^, Sgl^PRR1^, and Sgl^Hgal^) Sgls. Measurements of cells (n=25 for each Sgl) were done using the measurement tool from ImageJ. The box plot was created using GraphPad Prism 9. To minimize bias in the selection of cells with blebs for the analysis, all cells showing blebs in an image were measured.

**Figure 6.**
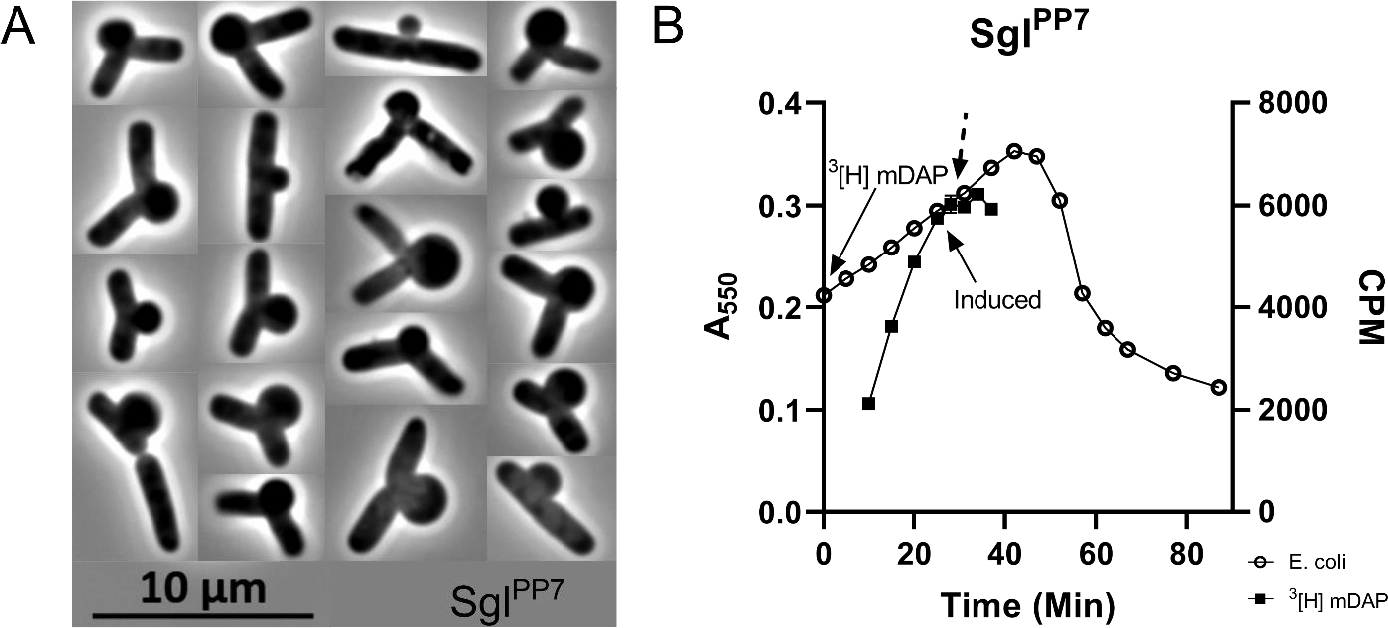
Sgl^PP7^ portrays type I lysis morphology and isotope labeling. **A**. Microscopy images of bleb formation after the induction of the Sgl^PP7^. **B**. Isotope labeling of the peptidoglycan layer using ^3^[H] mDAP. The open circles represent the optical density of the culture, and the closed squares represent the counts per minute of ^3^[H] mDAP incorporated into insoluble material. The solid arrow on the left indicates the time at which ^3^[H] mDAP was added to the growing culture. The middle solid arrow shows when the culture was induced with 0.4% L-arabinose and the dashed arrow points to the time at which incorporation is judged to have ceased.

### Type I character of the Sgl of Pseudomonas phage PP7

Among the Sgls originally assigned to the “L-like” or type II category by virtue of having a domain structure that resembled that established for L is the Sgl of *Pseudomonas* phage PP7, which uses the polar pilus for its receptor (8, 36). However, during this project, Adler and colleagues reported the results of a screen for multicopy suppressors for a number of Sgls; in this report, they identified *murJ* as a multicopy suppressor for PP7 (19). In the same work, this unexpected finding was supported by genetic analysis, including the isolation of missense mutations in *murJ* conferring resistance to Sgl^PP7^. In view of the apparent L-like domain structure, we wondered if Sgl^PP7^ might have both type I and type II character. However, applying the two methods developed here revealed solely type I phenotypes (Fig. 5, 6). Sgl^PP7^ caused an abrupt halt in net PG biosynthesis, as judged by ^3^[H]mDAP incorporation, as well as a septal bleb lysis morphology.

## Discussion

Here we show that the type I and type II Sgls can be distinguished by two methods, incorporation of ^3^[H]mDAP into PG and phase contrast microscopy of lysing cells. In addition, we provide physiological evidence that supports the recent finding that Sgl^PP7^ is a type I Sgl. The first criterion, incorporation of ^3^[H]mDAP into insoluble PG, is the most definitive and mechanistically relevant because the enzymes required for de novo biosynthesis and externalization of Lipid II, the immediate precursor of PG polymer, are known and highly conserved. With the discovery that Sgl^PP7^ is a MurJ inhibitor, there are now four different small lytic phages that have been shown to use a type I Sgl for host lysis: ϕX174 and Qβ targeting MraY and MurA, respectively and M and PP7 targeting MurJ. So far, none of the proteins in the multimeric complexes (i.e., the elongasome and the divisome; (31, 37) that incorporate the disaccharide subunit of Lipid II into the PG has shown up as an Sgl target. This could be interpreted as surprising, since the septal bleb morphological phenotype that characterizes type I Sgl function looks the same as that produced by beta-lactam antibiotics. However, efficient lysis in the context of a phage infection might require inhibition of both the elongasome and divisome complexes, which would necessitate that the Sgl target two different proteins, one in each complex. It is also worth noting that two of the four known type I Sgls target the same protein, MurJ, the Lipid II “flippase”. Moreover, all three of the type I Sgls that evolved within the boundaries of essential phage genes target membrane proteins (E targets MraY, and both Sgl^PP7^ and Sgl^M^ target MurJ). It may be easier to evolve a functional Sgl that is localized to the membrane, where the lipidic environment imposes helical structure on the polypeptide, than one required to fold in the aqueous phase. We have shown that, for candidate Sgls with TMDs, the ability to cause overt lysis in *E. coli* can be selected rapidly (20). In contrast, evolving de novo a polypeptide to target a soluble enzyme might be much easier with a pre-existing tertiary structural scaffold; i.e., A_2_ inhibiting MurA.

It should be noted that the original conclusion that L did not act by inhibiting cell wall synthesis used a pulse-labelling protocol and thus did not rule out the possibility that L acted by inducing PG degradation. Our results show that net PG synthesis is unaffected by L and therefore the balance between overall PG synthesis and degradation is unchanged. This does not rule out localized imbalances. The simplest general concept is that L interferes with proper distribution of PG biosynthesis, so that the multiprotein PG biosynthesis complexes are just as active but not positioned at the right places for homeostasis and growth. It is also important to note that at this point, type II Sgl function is mainly defined as the absence of type I characteristics. The notion that L was just one of many similarly functioning Sgls produced by convergent evolution was in part based on the attractive notion that the four-domain structure defined by genetic analysis of L seemed to be present in other Sgls. It is possible that Sgl^PP7^ has both Type I and Type II functionality, with the Type I characteristic manifesting a phenotypic dominance.

Finally, it must be noted that there is no evidence that these ssRNA genomes that are extracted from metatranscriptomes are from plaque-forming lytic phages. The ssRNA infection cycle does not directly interfere with the DNA-based life of the host cell, so it can be considered that many if not most RNA phages cause chronic infections in which virions accumulate indefinitely while allowing the host to continue growth and division cycles. Indeed, even with the known ssRNA phages, infections can lead to non-lytic intracellular accumulation of virions to levels of 10^4^ to 10^5^ per cell (38). Thus, many ssRNA phages may persist in an endemic carrier state; in this scenario, only occasional host lysis events occur every few generations, thus seeding the environment with virions that can take advantage of new host cells. The original isolate Q that gave rise to Qβ was just such a chronic phage, since it formed regions of retarded growth on plates, rather than sharply defined plaques and gave rise to bacterial colonies that constitutively produced virions (39). Modern Qβ likely arose from mutations that increased the affinity of A_2_ for MurA. In fact, single base changes that increase the production rate of A_2_ and overcome host resistance due to missense changes in MurA arise at a high rate in the context of the phage (40).

## Materials and Methods

### Bacterial strains, plasmids, and growth conditions

The bacterial strains used in this work unless otherwise stated were as follows DH5α (ThermoFisher Scientific). The plasmids and primers that were used in this study can be found in table 1. Bacterial cultures were grown in lysogeny broth (LB) with 100µg·mL^-1^ ampicillin at 37°C with constant aeration in a gyrotory water bath shaker (New Brunswick Scientific) for lysis curves and microscopy. Overnight cultures were grown in 18 × 150 mm glass culture tubes (Fisher Scientific 14-961-32). For overnight cultures of *E. coli* harboring the pBAD24 vector, glucose was added at a final concentration of 0.4% to prevent premature gene expression. For isotope labeling experiments, bacterial cultures were grown in defined M9 media supplemented with 0.2% glycerol, 0.01% thiamine, 100µg·mL^-1^ ampicillin, and L-amino acids (20 µgmL^-1^ His; 30 µg·mL^-1^ Gly, Ile, Phe, Pro, Ser, Val; 50 µg·mL^-1^ Arg; 60 µg·mL^-1^ Ala; 70 µg·mL^-1^ Asn, Lys; 80 µg·mL^-1^ Glu; 90 µg·mL^-1^ Gln; 100 µg·mL^-1^ Met, The; 110 µg·mL^-1^ Leu). High levels of Met and Thr were added to reduce the intracellular pool of mDAP (37). Induction of cultures was done with 0.4% (w/v) L-arabinose.

**Table 1.**
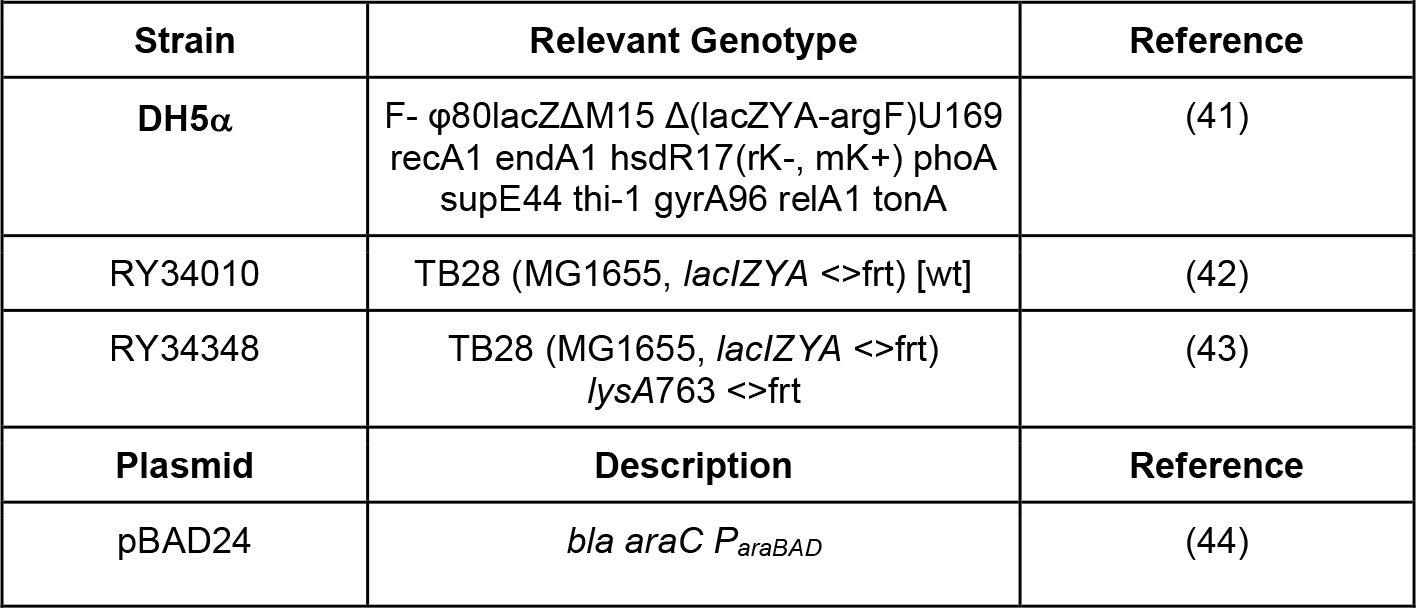
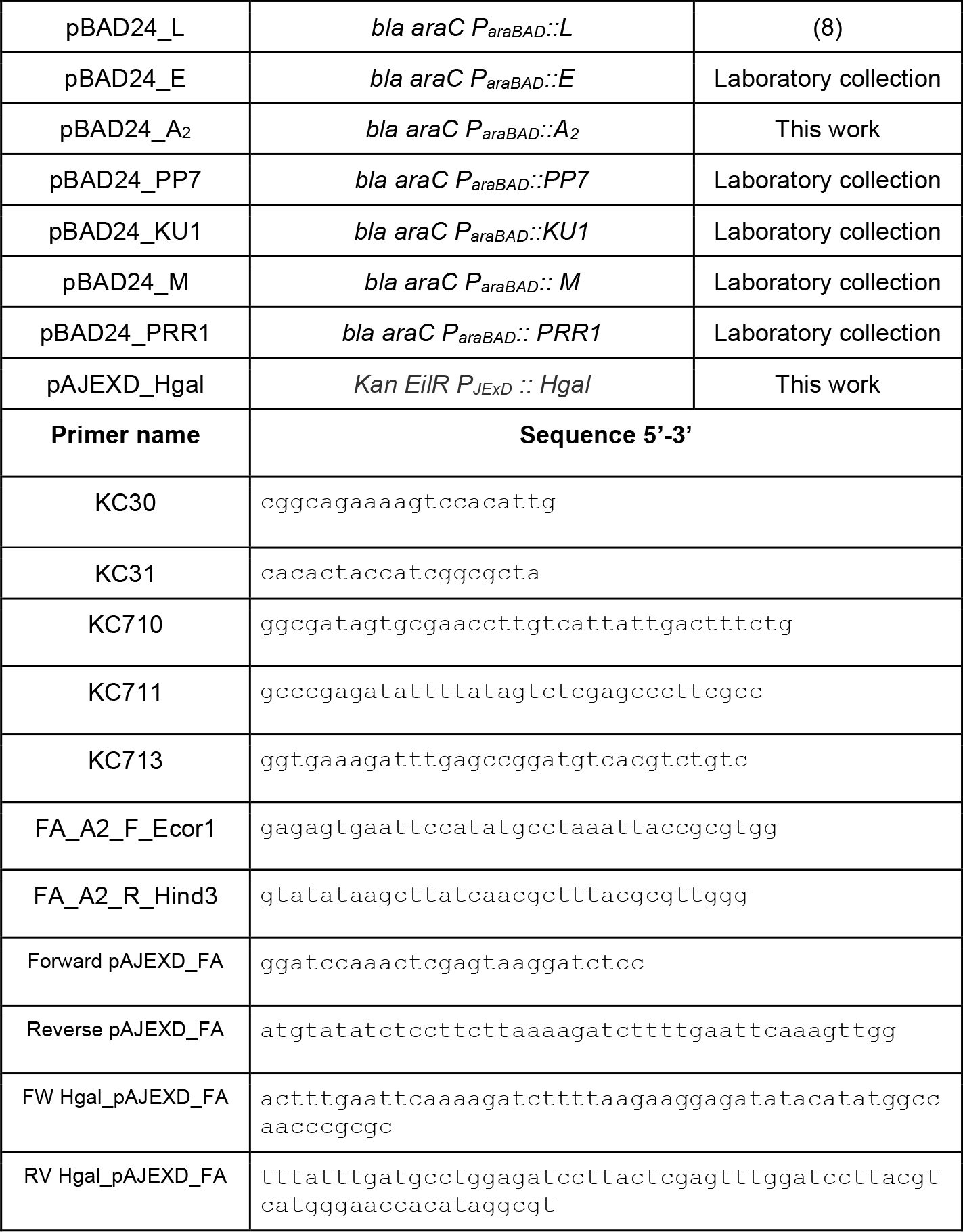

### Chemical Reagents and enzymes

Tritiated diaminopimelic (250µCi·mL^-1^) acid was purchased from VWR. All other chemicals were purchased from Sigma Aldrich unless otherwise stated.

### Microscopy

Prior to microscopy, lysis curves of each plasmid construct in strain RY34010 were done by inoculating a 250mL culture flask containing 25mL of LB and appropriate amount of antibiotic (100µg·mL^-1^ ampicillin or 40µg·mL^-1^ kanamycin) with a 1:200 dilution of bacteria. Cultures were grown with aeration in a water bath until they reached an A_550_ of ~0.2, at which they were then induced with 0.4% L-arabinose or 0.075µM of crystal violet. For all microscopy images, samples were taken ten minutes prior, five minutes prior, and at the time of expected lysis .At each time point 1.5µL samples were put onto a 25 × 75 x 1mm glass slide (Globe Scientific) and covered with a 22 × 22mm cover slip (Globe Scientific) and imaged in inverted orientation at 100X with oil immersion on a Zeiss microscope. Slides were imaged at an exposure of 100ms for no longer than 20 minutes at a time to avoid artifacts.

### Image Analysis

The images were analyzed using ImageJ (45). The length from each pole to the center of the bleb was measured three times. The center of the bleb can be defined as the region at the base of the bleb that is equidistant from the left and right side of the bleb (protruding membranes). These measurements were used to calculate the total length of the cell as well as calculate the location of the bleb.

### ^3^[H]mDAP incorporation and measurement of PG synthesis

For the labeling experiments, strains were grown in 25mL of defined media in 250mL culture flasks (mother culture) until an A_550_ ~0.1 was reached. At this point, 5mL of the mother culture was placed into a prewarmed 50mL culture flask (daughter culture). Both cultures were then left to grow to an A_550_ ~0.2; growth of cultures was measured via the mother flask. Once the culture reached the desired optical density (OD) ^3^[H]mDAP was added to the daughter flask to a final level of 1µCi·mL^-1^ and given a 10-minute pre-labeling period. After the pre-labeling period, 200µL samples were taken at designated time points (Fig. 3) and put into a microcentrifuge tube with 800µL boiling 5% SDS and vortexed vigorously. The samples were boiled for an hour and a half and then cooled to room temperature. After cooling to room temperature, the samples were vortexed vigorously again and briefly spun down. Filters were rinsed with 5mL of lysine (70 µg·mL^-1^) before the addition of any sample. After the blocking step 200µL of each sample was filtered through a 0.22µm cellulose nitrate membrane filter (Cytiva), washed with 30mL of the lysine-water solution, and left to dry overnight. For type I Sgls each time point had two samples filtered. Such that an average for each time point could be taken. Next the filters were put into plastic 20mL vials (Fisher Scientific) with Ecoscint A scintillation fluid (National Diagnostics). The vials with the filters were then kept in the dark for a minimum of 1 hour prior to reading. The counts per minute of the samples were read in a Tri-Carb 2910 TR liquid scintillation counter (PerkinElmer) with a count time of 1 minute.

## Acknowledgements

The authors want to thank Diobenhi Castellanos for her assistance with the lysis curve replicates. We also thank Jennifer Tran and Lorna Min for assistance in cloning and for preliminary experiments in cell morphology.

## Contributions

S.F.A, and R.Y. were primarily responsible for the design of the studies and analysis of the results. T.G.B. designed and executed the original radioactive labeling experiments. S.F.A. performed most of the physiological experiments reported and s and created the figures. S.F.A and R.Y prepared and edited the manuscript. K.C. and T.G.B provided edits on the manuscript.

## References

1. Cahill J, Young R. 2019. Phage Lysis: Multiple Genes for Multiple Barriers, p 33–70, Advances in Virus Research. Elsevier.

2. Young R. 2014. Phage lysis: Three steps, three choices, one outcome. Journal of Microbiology 52:243–258.

3. Chamakura K, Young R. 2019. Phage single-gene lysis: Finding the weak spot in the bacterial cell wall. Journal of Biological Chemistry 294:3350–3358.

4. Bernhardt TG, Wang IN, Struck DK, Young R. 2002. Breaking free: “protein antibiotics” and phage lysis. Res Microbiol 153:493–501.

5. Fischetti VA. 2008. Bacteriophage lysins as effective antibacterials. Current Opinion in Microbiology 11:393–400.

6. Atkins JF, Steitz JA, Anderson CW, Model P. 1979. Binding of mammalian ribosomes to MS2 phage RNA reveals an overlapping gene encoding a lysis function. Cell 18:247–56.

7. Berkhout B, De Smit MH, Spanjaard RA, Blom T, Van Duin J. 1985. The amino terminal half of the MS2-coded lysis protein is dispensable for function: implications for our understanding of coding region overlaps. The EMBO Journal 4:3315–3320.

8. Chamakura KR, Edwards GB, Young R. 2017. Mutational analysis of the MS2 lysis protein L. Microbiology 163:961–969.

9. Chamakura KR, Tran JS, Young R. 2017. MS2 Lysis of Escherichia coli Depends on Host Chaperone DnaJ. Journal of Bacteriology 199:JB.00058-17.

10. Reed CA, Langlais C, Kuznetsov V, Young R. 2012. Inhibitory mechanism of the Qβ lysis protein A2. Mol Microbiol 86:836–44.

11. Chamakura KR, Sham L-T, Davis RM, Min L, Cho H, Ruiz N, Bernhardt TG, Young R. 2017. A viral protein antibiotic inhibits lipid II flippase activity. Nature Microbiology 2:1480–1484.

12. Vollmer W, Blanot D, de Pedro MA. 2008. Peptidoglycan structure and architecture. FEMS Microbiol Rev 32:149–67.

13. Silhavy TJ, Kahne D, Walker S. 2010. The Bacterial Cell Envelope. Cold Spring Harbor Perspectives in Biology 2:a000414–a000414.

14. Bernhardt TG, Wang IN, Struck DK, Young R. 2001. A protein antibiotic in the phage Qbeta virion: diversity in lysis targets. Science 292:2326–9.

15. Holtje J, van Duin J. 1984. MS2 phage induced lysis of E. coli depends upon the activity of the bacterial autolysins. Elsevier Science Publishers, New York, NY.

16. Walderich B, Höltje JV. 1989. Specific localization of the lysis protein of bacteriophage MS2 in membrane adhesion sites of Escherichia coli. J Bacteriol 171:3331–6.

17. Goessens WH, Driessen AJ, Wilschut J, Van Duin J. 1988. A synthetic peptide corresponding to the C-terminal 25 residues of phage MS2 coded lysis protein dissipates the protonmotive force in Escherichia coli membrane vesicles by generating hydrophilic pores. The EMBO Journal 7:867–873.

18. Julija M, Janosch M, Clara B, Volker D, Achilleas Stefanos F, Nina M, Frank B. 2023. In vitro characterization of the phage lysis protein MS2-L. In vitro characterization of the phage lysis protein MS2-L 2:28.

19. Adler BA, Chamakura K, Carion H, Krog J, Deutschbauer AM, Young RF, Mutalik VK, Arkin AP. 2022. Parallel multicopy-suppressor screens reveal convergent evolution of phage-encoded single gene lysis proteins. Cold Spring Harbor Laboratory.

20. Chamakura KR, Tran JS, O’Leary C, Lisciandro HG, Antillon SF, Garza KD, Tran E, Min L, Young R. 2020. Rapid de novo evolution of lysis genes in single-stranded RNA phages. Nat Commun 11:6009.

21. Starr EP, Nuccio EE, Pett-Ridge J, Banfield JF, Firestone MK. 2019. Metatranscriptomic reconstruction reveals RNA viruses with the potential to shape carbon cycling in soil. Proc Natl Acad Sci U S A 116:25900–25908.

22. Callanan J, Stockdale SR, Shkoporov A, Draper LA, Ross RP, Hill C. 2020. Expansion of known ssRNA phage genomes: From tens to over a thousand. Sci Adv 6:eaay5981.

23. Neri U, Wolf YI, Roux S, Camargo AP, Lee B, Kazlauskas D, Chen IM, Ivanova N, Allen LZ, Paez-Espino D, Bryant DA, Bhaya D, Krupovic M, Dolja VV, Kyrpides NC, Koonin EV, Gophna U. 2022. A five-fold expansion of the global RNA virome reveals multiple new clades of RNA bacteriophages. Cold Spring Harbor Laboratory.

24. Domingo E, Holland JJ. 1997. RNA VIRUS MUTATIONS AND FITNESS FOR SURVIVAL. Annual Review of Microbiology 51:151–178.

25. Bernhardt TG, Struck DK, Young R. 2001. The Lysis Protein E of φX174 Is a Specific Inhibitor of the MraY-catalyzed Step in Peptidoglycan Synthesis. Journal of Biological Chemistry 276:6093–6097.

26. Paliy O, Gunasekera TS. 2007. Growth of E. coli BL21 in minimal media with different gluconeogenic carbon sources and salt contents. Applied Microbiology and Biotechnology 73:1169–1172.

27. Thakur CS, Brown ME, Sama JN, Jackson ME, Dayie TK. 2010. Growth of wildtype and mutant E. coli strains in minimal media for optimal production of nucleic acids for preparing labeled nucleotides. Applied Microbiology and Biotechnology 88:771–779.

28. Bremer H, Churchward G, Young R. 1979. Relation between growth and replication in bacteria. J Theor Biol 81:533–45.

29. Lederberg J, St Clair J. 1958. Protoplasts and L-type growth of Escherichia coli. J Bacteriol 75:143–60.

30. Ciak J, Hahn FE. 1957. Penicillin-induced lysis of Escherichia coli. Science 125:119–20.

31. Rohs PDA, Bernhardt TG. 2021. Growth and Division of the Peptidoglycan Matrix. Annu Rev Microbiol 75:315–336.

32. Bradley DE, Dewar CA, Robertson D. 1969. Structural Changes in Escherichia coli Infected with a X174 Type Bacteriophage. Journal of General Virology 5:113–121.

33. Henrich B, Lubitz W, Plapp R. 1982. Lysis of Escherichia coli by induction of cloned ϕX174 genes. Molecular and General Genetics MGG 185:493–497.

34. Young KD, Young R. 1982. Lytic action of cloned phi X174 gene E. Journal of Virology 44:993–1002.

35. Blasi U, Henrich B, Lubitz W. 1985. Lysis of Escherichia coli by Cloned X174 Gene E Depends on its Expression. Microbiology 131:1107–1114.

36. Brockhurst MA, Buckling A, Rainey PB. 2005. The effect of a bacteriophage on diversification of the opportunistic bacterial pathogen, Pseudomonas aeruginosa. Proc Biol Sci 272:1385–91.

37. Egan AJF, Errington J, Vollmer W. 2020. Regulation of peptidoglycan synthesis and remodelling. Nat Rev Microbiol 18:446–460.

38. Bradley DE. 1966. The Structure and Infective Process of a Pseudomonas Aeruginosa Bacteriophage Containing Ribonucleic Acid. Microbiology 45:83–96.

39. Miyake T, Yanagisawa K, Watanabe I. 1966. Alteration of a host character by infection with RNA phage Q-beta. Jpn J Microbiol 10:141–8.

40. Reed CA, Langlais C, Wang IN, Young R. 2013. A(2) expression and assembly regulates lysis in Qβ infections. Microbiology (Reading) 159:507–514.

41. Jones PA, Samuels NM, Phillips NJ, Munson RS, Jr., Bozue JA, Arseneau JA, Nichols WA, Zaleski A, Gibson BW, Apicella MA. 2002. Haemophilus influenzae type b strain A2 has multiple sialyltransferases involved in lipooligosaccharide sialylation. J Biol Chem 277:14598–611.

42. Bernhardt TG, De Boer PAJ. 2003. The Escherichia coli amidase AmiC is a periplasmic septal ring component exported via the twin-arginine transport pathway. Molecular Microbiology 48:1171–1182.

43. Cho H, Uehara T, Bernhardt G, Thomas. 2014. Beta-Lactam Antibiotics Induce a Lethal Malfunctioning of the Bacterial Cell Wall Synthesis Machinery. Cell 159:1300–1311.

44. Guzman LM, Belin D, Carson MJ, Beckwith J. 1995. Tight regulation, modulation, and high-level expression by vectors containing the arabinose PBAD promoter. J Bacteriol 177:4121–30.

45. Schneider CA, Rasband WS, Eliceiri KW. 2012. NIH Image to ImageJ: 25 years of image analysis. Nat Methods 9:671–5.

