## Supplemental Figures 1-4 for "Physiological characterization of single gene lysis proteins"

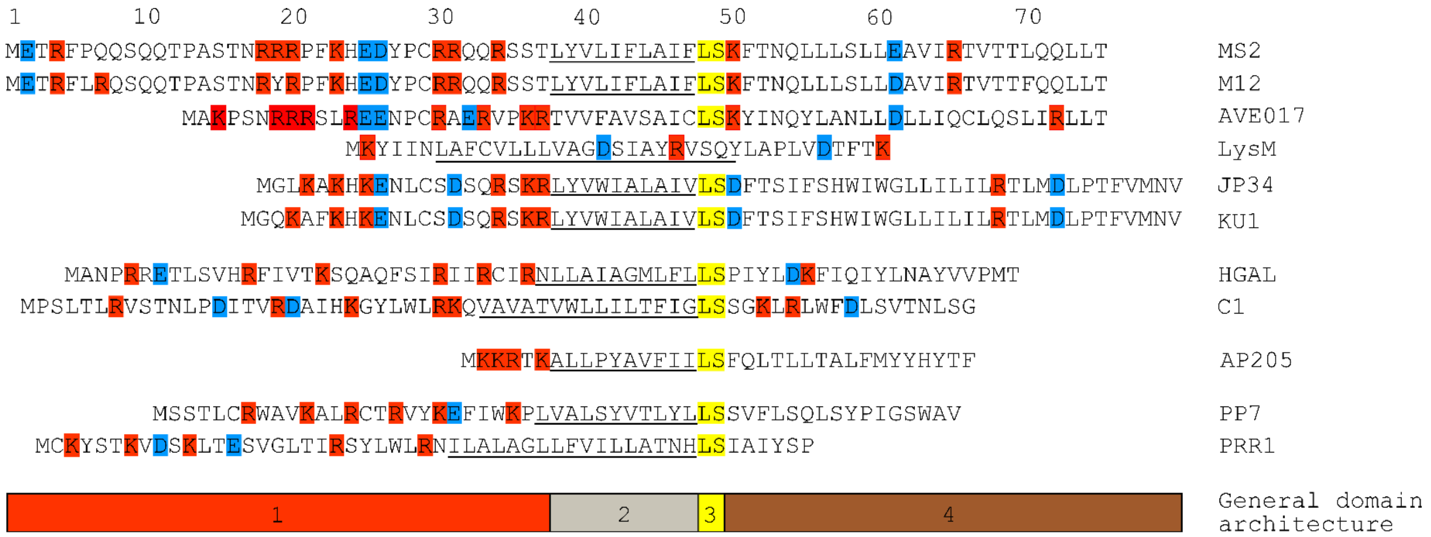


**Supplementary Figure 1. MS2-like Sgl sequence alignment**

Alignment of the proposed L-like Sgls from F-specific phages to the L protein from canonical ssRNA phage MS2. GenBank files for the Sgls shown from top to bottom are: MS2 (CAA23990.1), M12 (AAF19634.1), AVE017 (https://doi.org/10.1371/journal.pbio.1002409.s001), LysM (YP_007111576.1), JP34 (AAA72211.1), KU1 (AAF67675.1), HGAL (YP_007237174.1), C1 (YP_007237128.1), AP205 (NP_085469.1), PP7 (NP_042306.1), PRR1 (YP_717670.1). The schematic of the general domain architecture at the bottom is as follows: 1. basic N-terminus 2. hydrophobic domain 3. leucine-serine motif 4. C-terminus. Adjusted from (8)

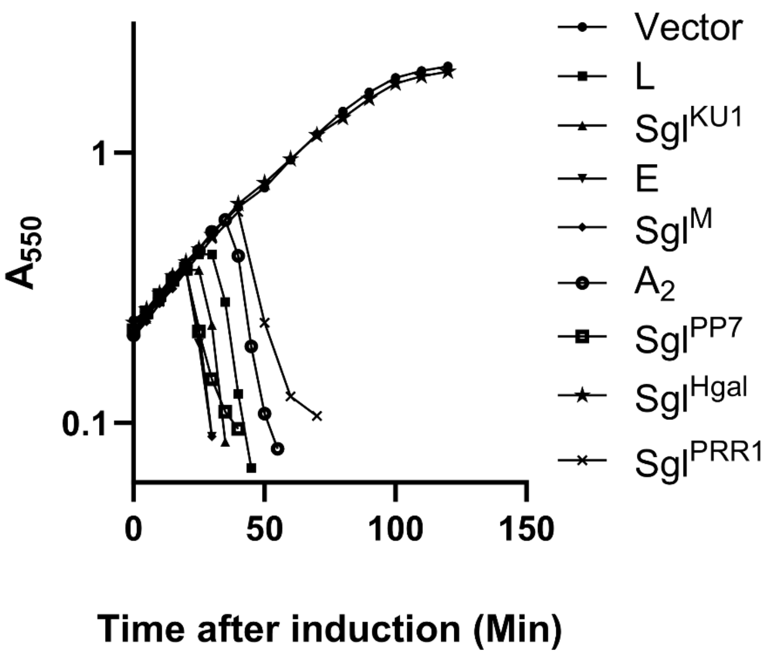


**Supplementary Figure 2. Lysis profiles for *Sgls***

Cultures of *E. coli* harboring different Sgls were grown in lysogeny broth at 37℃ with aeration and induced with L-arabinose once the $OD_{550}$ reached ~0.2. From top to bottom: Vector (closed circles), L (closed squares), $Sgl^{KU1}$ (closed upward triangle), E (closed downward triangle), $Sgl^{M}$ (closed diamond), $A_2$ (open circles), $Sgl^{PP7}$ (open squares), $Sgl^{Hgal}$ (stars), and $Sgl^{PRR1}$ (X's). The lysis curve is representative of three biological replicates.

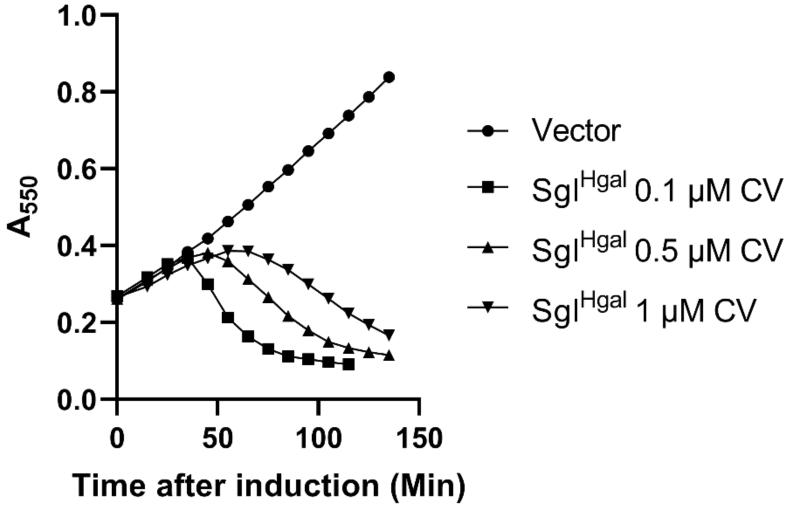


**Supplementary Figure 3. Titration lysis profile for Hgal**

From top to bottom: pAJEXD vector with no Sgl (closed circles), $Sgl^{Hgal}$ induced with 0.1µM CV (closed squares), $Sgl^{Hgal}$ induced with 0.5µM CV (closed upward triangles), $Sgl^{Hgal}$ induced with 1 µM CV (closed downward triangles). The lysis curves are representative of three biological replicates.

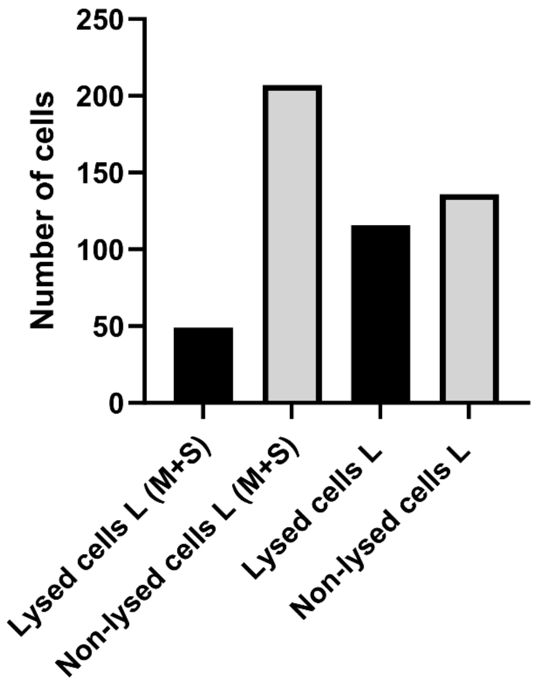


**Supplementary Figure 4. Lysis frequency of *E. coli* from induction of L with and without stabilizing agents**

Cultures of *E. coli* harboring the Sgl L were grown in LB with and without 10mM $MgCl_2$ and 0.2M Sucrose (M+S) at 37℃ to an optical density of ~0.2 and then induced with 0.4% L-arabinose. Ten minutes prior to lysis, 1.5μL samples were taken from culture flasks, put onto glass slides, covered with a cover slip, and imaged using the 40X objective lens. ~250 cells were counted for stabilizing conditions (M+S) and non-stabilizing conditions.
